# Rheo-2DIR spectroscopy reveals strain-induced hydrogen-bond redistribution in polyurethane

**DOI:** 10.1101/2022.10.04.510759

**Authors:** Giulia Giubertoni, Michiel Hilbers, Hajo Groen, Anne Van der Weide, Daniel Bonn, Sander Woutersen

**Affiliations:** University of Amsterdam, Science Park 904, 1098XH Amsterdam, The Netherlands

## Abstract

The remarkable elastic properties of polymers are ultimately due to their molecular structure, but the relation between the macroscopic and molecular properties is often difficult to establish, in particular for (bio)polymers that contain hydrogen bonds, which can easily rearrange upon mechanical deformation. Here we show that two-dimensional infrared spectroscopy on polymer films in a miniature stress tester sheds new light on how the hydrogen-bond structure of a polymer is related to its visco-elastic response. We study thermoplastic polyurethane, a block copolymer consisting of hard segments of hydrogen-bonded urethane groups embedded in a soft matrix of polyether chains. The conventional infrared spectrum shows that upon deformation, the number of hydrogen bonds increases, a process that is largely reversible. However, the 2DIR spectrum reveals that the distribution hydrogen-bond strengths becomes slightly narrower after a deformation cycle, due to the disruption of weak hydrogen bonds, an effect that could explain the strain-cycle induced softening (Mullins effect) of polyurethane. These results show how rheo-2DIR spectroscopy can bridge the gap between the molecular structure and the macroscopic elastic properties of (bio)polymers.

---

The unique strength and resilience of polymeric materials such as plastics and biopolymer networks finds its origin in the molecular-scale structural changes that the polymer chains undergo when subjected to deformation. A detailed understanding of the connection between the macroscopic and microscopic properties of polymers is essential, not only for predicting the mechanical properties of synthetic polymers, but also for understanding the molecular origin of dysfunctional polymer systems, such as occur in for instance collagen-related diseases. The most straightforward way to investigate the molecular origin of polymer elastic response is to directly observe the changes in molecular structure induced by externally applied strain. Such experiments have used a range different structural probing methods, notably X-ray diffraction,^1–3^ but also Raman and infrared (IR) spectroscopy. ^4–10^

In the case of hydrogen-bonded polymer networks, the strain-induced structural changes generally involve rearrangement of the hydrogen bonds between molecular groups of adjacent (bio)polymer chains. Among the above-mentioned techniques, IR spectroscopy is very suitable to investigate such rearrangements, since the frequencies and line shapes of the vibrational transitions contain detailed information on the hydrogen-bond structure. ^11^ Combining rheology with infrared spectroscopy (rheo-IR) to study the molecular changes in polymers under applied strain (Fig. 1A) has provided a detailed molecular picture of the strain-induced molecular rearrangements in a broad range of polymers.^4–9^ However, infrared absorption spectra are often rather congested, and it can be difficult to disentangle overlapping vibrational bands; and although the absorption frequency is often a good indicator of the local structure and/or environment, it is generally difficult to derive unambiguous conclusions from the frequencies alone, since they are determined by more than one effect (e.g. the conformation and solvent interactions). Furthermore, with conventional IR spectroscopy, inhomogeneous spectral broadening (due to a distribution of transition frequencies) cannot be easily separated from homogeneous spectral broadening (caused by fast frequency fluctuations of individual vibrations).

**Figure 1:**
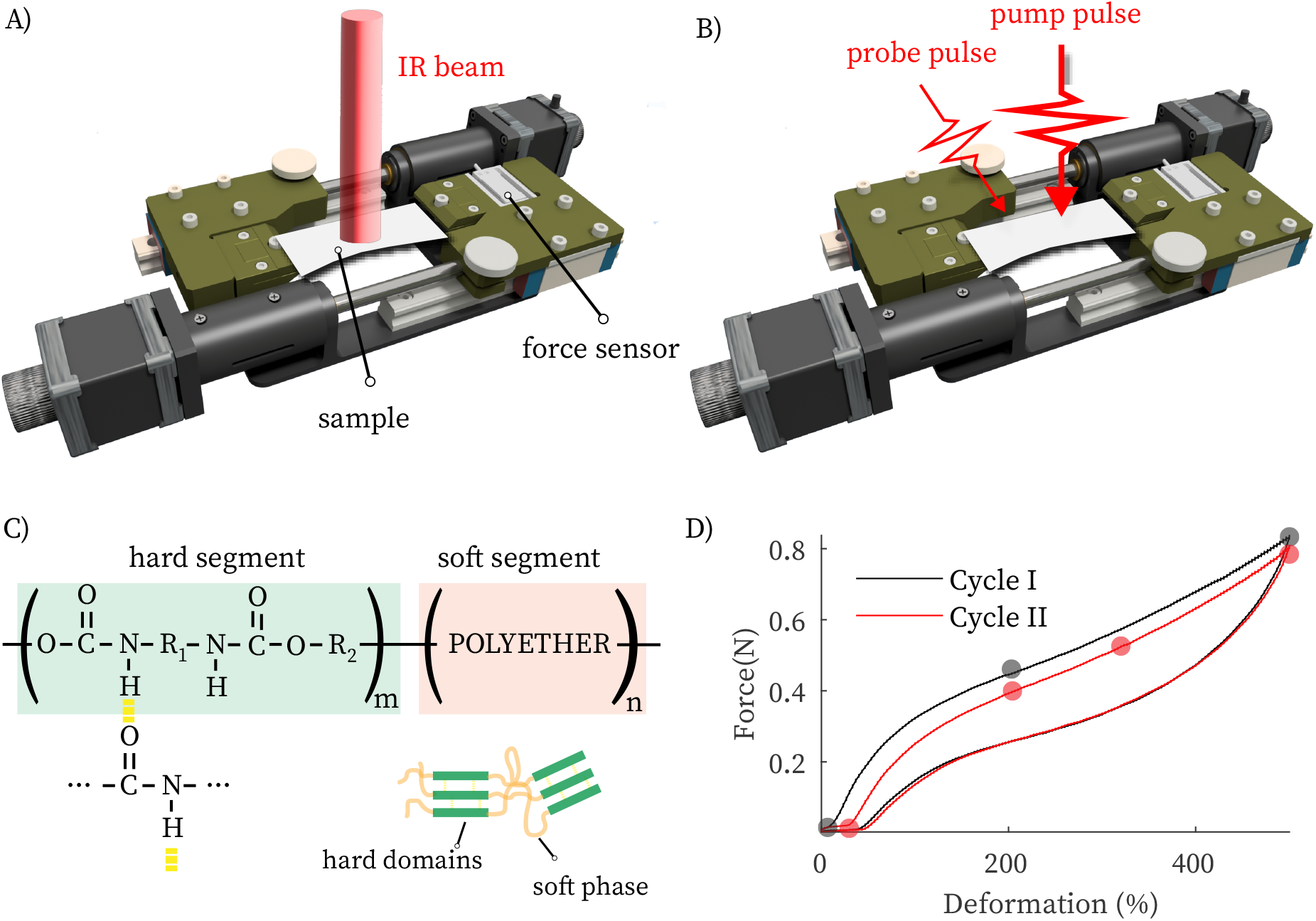
(A, B) Schematic of rheo-IR and rheo-2DIR. (C) Structure of polyurethane, a block co-polymer composed of soft and hard segments. The urethane groups in the hard segments form strong hydrogen bonds that act as physical cross links. Because of the different polarity and chemical nature, the soft and hard segments separate, leading to the formation of hard domains embedded in a soft phase. D) Stress-strain curve of a TPU film at a strain rate of 8 s with a strain step-size of 3%. The stress–strain curve in the second cycle is more compliant than in the first cycle (strain softening). Filled circles indicate the deformations at which rheo-2DIR was performed.

These challenges can be addressed by means of two-dimensional infrared (2DIR) spectroscopy, which makes it possible to separate overlapping spectral bands, measure vibrational couplings, and separate the homogeneous and inhomogeneous contributions to the line broadening.^12^ Inspired by rheo-IR spectroscopy, we here combine two-dimensional infrared spectroscopy with rheometry by inserting a miniature universal stress tester into a 2DIR setup (rheo-2DIR). We use this new method to investigate strain-induced changes in the hydrogen-bond distribution of thermal polyurethane, one of the most commonly used polymers, that exhibits strain-cycle induced softening behavior that is believed to be related to changes in the hydrogen-bond structure.^2^

A detailed description of the 2DIR setup can be found in refs. 13 and 14. Briefly, we use an amplified Ti:sapphire laser combined with an optical parametric amplifier and difference-frequency generation to generate tunable mid-IR pulses (∼20 *µ*J, ∼6100 nm) with a spectral width of 150 cm^−1^ (FWHM) at 1 kHz repetition rate. The IR beam is split into a probe and reference beam (each 5%), and a pump beam (90%) that is aligned through a Fabry-Pérot interferometer. The pump and probe beams are overlapped in the sample (∼250 *µ*m focal diameter), and the transmitted spectra of the probe pulse in the presence and absence of the pump pulse are recorded with a 32-pixel mercury cadmium telluride array. The pump and probe polarizations are at the magic angle (54.5^*◦*^) to obtain polarization-independent spectra. ^12^ In the rheo-2DIR setup (Fig. 1B), the polymer films are clamped on both sides in a miniature home-built stress tester inserted in the 2DIR setup, and controlled deformations are applied by moving the two clamps in opposite directions with 2 *µ*m precision, using steppermotors (Physik Instrumente). The force is measured with a precision of 0.001 N using a force sensor (KD34s, ME-Systeme) positioned on one of the clamps. Polyurethane films were purchased in the form of polyurethane condoms from Protex (France) and Sagami (Japan). To remove the lubricant, the samples were dried and cleaned with ethanol. The film thickness was 31 ± 5 *µ*m (Protex 002) and 18 ± 5 *µ*m (Sagami 001).

Thermoplastic polyurethane (TPU) is a block co-polymer composed of urethane- and polyether-based segments (Fig. 1C). At room temperature, the polyether (“soft”) domains are above their glass transition temperature, and give TPU its rubber-like behavior; the polyurethane (“hard”) domains are below their glass or melt transition temperature and are believed to give rise to the hysteresis, permanent deformation, high modulus, and tensile strength. ^15^ The mechanical response to deformation of TPU involves changes in the arrangement and strength of the hydrogen-bonds formed between the urethane links within hard segments and between the urethane links of hard and soft segments. In particular the stress-softening behavior upon recovery of zero-stress condition after deformation (Mullins effect) is believed to be connected to the disruption of weak hydrogen-bonds between hard and soft domains.^2^

Figure 1D shows stress-strain measurements where we deform a thin polyurethane film up to a final strain of 500%. We observe an elastically rigid response up to 100% (elastic regime), and then a more compliant response at higher strain (strain softening), until around 400% where the polyurethane again stiffens (strain hardening) as it approaches the fracture point. During unloading, we observe a large hysteresis and a significant residual strain. The mechanical response of the sample during the second cycle is much more compliant compared to the first cycle. This strain-softening behavior (the Mullins effect) ^15^ has been extensively investigated, but its molecular origins are still not completely understood. ^1,2^ Here, we show that rheo-2DIR spectroscopy can shed new light on this phenomenon by revealing how the hydrogen-bond distribution changes upon deformation.

Figure 2A shows the conventional IR spectrum of the polyurethane film (Sagami 001) in the carbonyl-stretch region. We observe two intense bands at 1703 and 1733 cm^−1^, which are due to the hydrogen-bonded (1703 cm^−1^) and free (1733 cm^−1^) carbonyl groups. ^16–20^ The lowering of the CO-stretch frequency upon hydrogen-bond formation is a well-known effect, that can be used as a sensitive probe of the hydrogen-bond structure. The peak of the hydrogen-bonded carbonyl groups is much broader than that of the free CO groups, and its shape reflects the distribution of hydrogen-bond strengths in the sample (convoluted with the homogeneous lineshape). Fig. 2B shows IR spectra of polyurethane film at different strains (the spectra have been normalized to the spectral area to correct for sample thinning due to stretching). Going from 0 to 200% strain, the intensity of the free-carbonyl peak decreases, while the intensity of the hydrogen-bonded carbonyl increases. At higher (500%) deformation, the hydrogen-bonded carbonyl peak slightly decreases in intensity, and broadens, while the free carbonyl does not show a significant change with respect to 200%. This is more clearly visible in Fig. 2C, where we plot the normalized intensity as a function of frequency and strain. We observe that the intensity of the hydrogen-bonded carbonyl increases when TPU enters the strain-softening regime, while it decreases when approaching the strain-hardening regime. In the strain-softening regime, stress is relieved by the conformational rearrangement of the chains, leading to a strain-induced ordering in the network. The increase in the number of hydrogen bonds at moderate strain reflects this increased ordering because the enhanced alignment of the polyurethane chains facilitates hydrogen-bond formation between urethane groups (this behavior is similar to strain-induced crystallization in natural rubber^18,19,21,22^). At higher strain, the finite extensibility of the chains leads to an upturn of the stress-strain curve. When approaching the maximum extension of the chain, hydrogen bonds are weakened and broken, leading to a decrease in intensity of the hydrogen-bonded carbonyl vibration in the strain-hardening region. ^23^ To summarize, rheo-IR shows that in the strain-softening regime, the number of hydrogen-bond increases because of an enhanced alignment of the chains, while it decreases in the strain-hardening regime because hydrogen bonds are weakened and broken while resisting extension.

**Figure 2:**
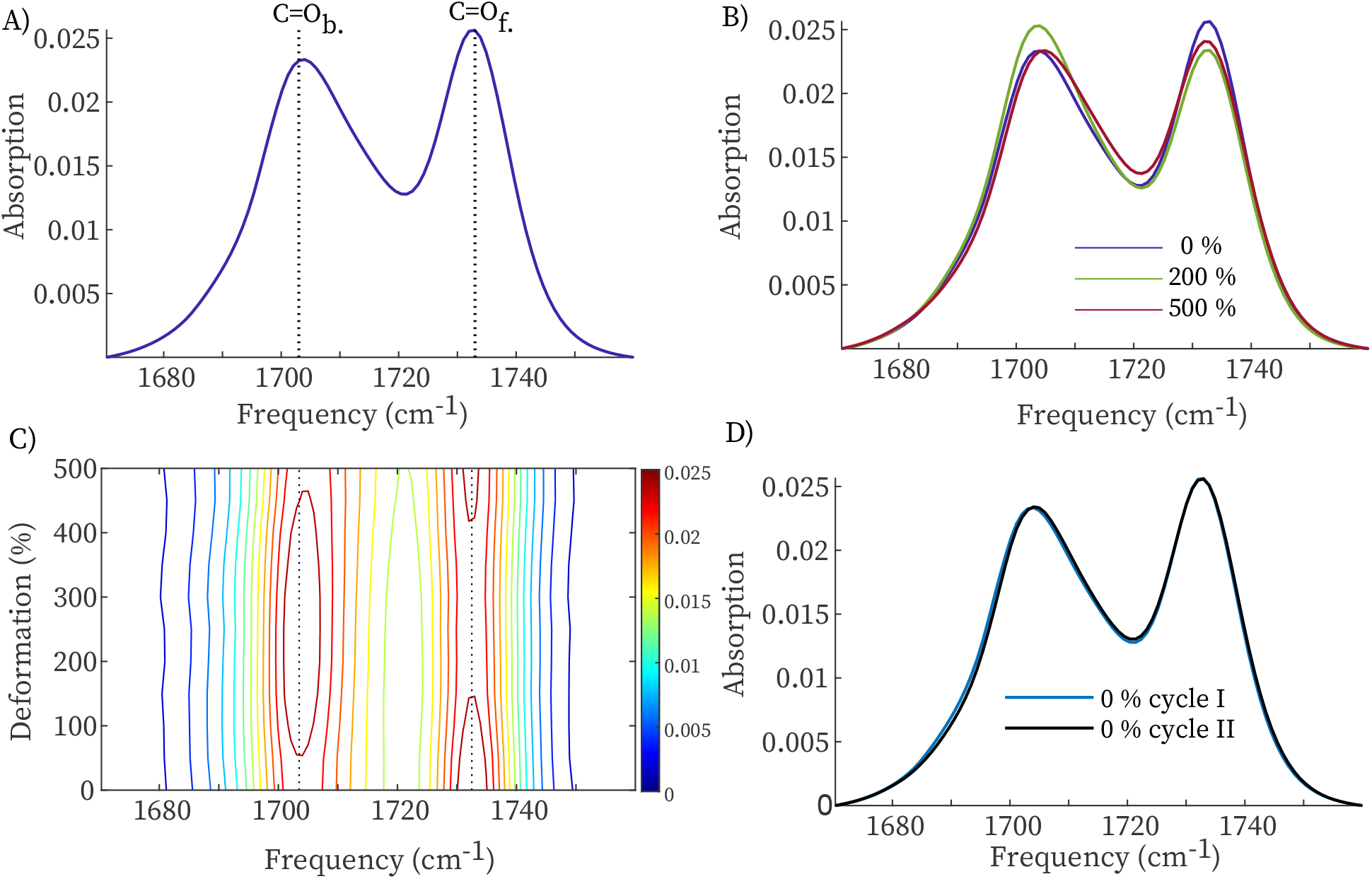
A) IR absorption spectra of TPU film at 0 % deformation. B) IR spectra of TPU at 0, 200, 500 % deformation normalized with respect to the respective total spectrum area to compensate for thinning effect. C) 2D gradient FT-IR map with respect to the deformation percentage. D) IR spectra of before deformation and upon recovery of the zerostress condition.

After unloading the sample, the infrared spectrum is very similar to the one observed before the deformation cycle (Fig. 2D) with the relative intensity between the two carbonyl bands returning to its value before deformation. This is somewhat surprising given that the macroscopic elastic properties have changed significantly (Fig. 1D). However, subtle changes in the vibrational lineshapes are generally difficult to observe in the linear IR spectrum, because the wings of the absorption bands are difficult to distinguish from the background absorption; and for strained polymer samples the situation is worsened since the deformation cycle causes changes in the background spectrum. To investigate strain-induced changes in the lineshape in detail, we therefore use 2DIR spectroscopy. Two important advantages of 2DIR with respect to conventional IR spectroscopy are (1) the homogeneous and inhomogeneous contributions to the lineshape are observed separately, and (2) the background absorption does not contribute to the 2DIR signal (since the 2DIR response scales as *µ*^4^, while conventional IR scales as *µ*^2^, where *µ* is the transition dipole moment of the vibrational transition).^12^ Figure 3A shows the 2DIR spectrum of the hydrogen-bonded carbonyl groups at 0% strain. In pump-probe 2DIR spectroscopy, we use a tunable narrow-band (10 cm^−1^ FWHM) pump pulse to excite molecular vibrations (in this case the CO-stretch vibration) at a specific frequency *ν*_pump_, and measure the pump-induced change in absorption Δ*A* at all frequencies using a broad-band probing pulse that is detected in frequency-resolved manner. Plotting Δ*A*(*ν*_probe_, *ν*_pump_) we obtain two-dimensional IR spectra. ^12^ In Figure 3A, positive Δ*A* is plotted as red contour areas, and negative Δ*A* as blue contour areas.

**Figure 3:**
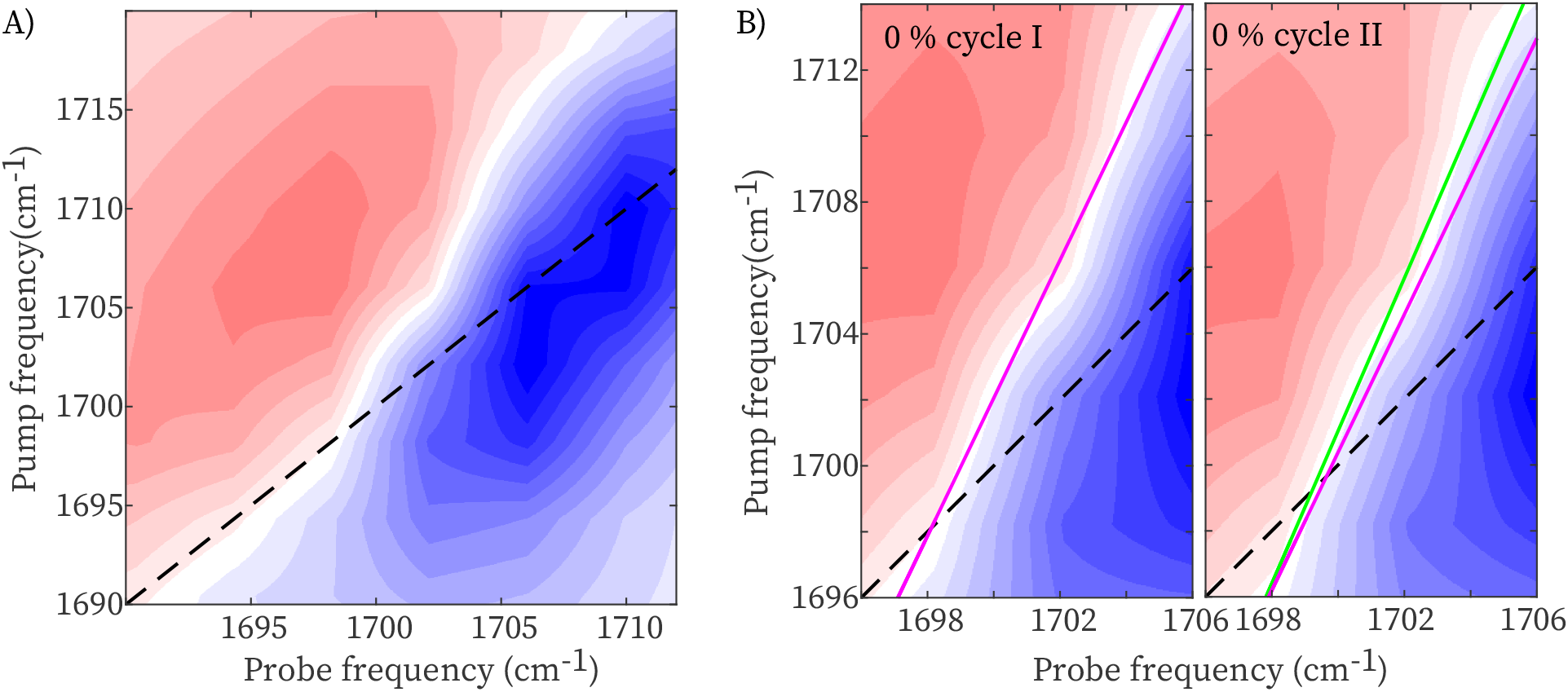
(A) 2DIR spectrum of TPU at zero stress condition. (B) 2DIR spectra and nodal line slopes at zero stress and upon recovery of zero-stress condition after a deformation up to 500%. The nodal line slope at zero stress is shifted along the probe axis in the right 2DIR spectrum for comparison.

The resonant excitation of the *v* = 0 → 1 transition by the pump pulse causes a decrease in absorption at the *v* = 0 → 1 transition (due to depletion of the *v* = 0 state and *v* = 1 → 0 stimulated emission) and an increase in absorption at the *v* = 1 → 2 transition frequency. ^12^ The dependence of the 2DIR response on the pump frequency is a measure of the inhomogeneous broadening of the IR band, and the two-dimensional lineshape can be used to disentangle inhomogeneous and homogeneous contributions to the line broadening.^12^ Since the inhomogeneous lineshape reflects the distribution of hydrogen-bond strengths, we can thus investigate changes in this distribution caused by the strain cycle. The simplest and most robust parameter to characterize the extent of inhomogeneous broadening is the inverse value of the slope (“nodal line slope”) of the 2DIR contours: ^24,25^ in the limiting case of purely homogeneous broadening, the Δ*A* contours are vertically aligned (zero nodal line slope, no dependence of the response on the pump frequency except for overall amplitude), whereas in the case of purely inhomogeneous broadening the slope is 1. In Figure 3B we compare the 2DIR spectra at zero stress and upon recovery of zero-stress after a deformation up to 500%, where we indicate the nodal line slopes as determined from global least-squares fits (Figure 4A shows the value during the deformation cycle). After the deformation cycle, the nodal line slope has decreased, indicating a decrease in the inhomogeneous width. To be certain that the observed subtle change in slope is significant, we have repeated the experiments several times, and on different TPU samples (Fig. 4B). The initial values of the nodal line slopes show small variations (probably due to variation in polymer composition across the condoms), but the decrease of the slope after the deformation cycle is reproducible. Repeating the deformation cycle on the same sample does not further change the slope (see SI), similar to the stress-strain curve, which also tends to stabilize, with most of the softening occurring in the first deformation cycle. The decrease of the nodal line slope indicates that the hydrogen-bond distribution has become narrower after a deformation cycle, suggesting that part of the hydrogen-bonds, which are broken during the loading, are not reformed upon recovery of zero-stress condition. This is confirmed by Fig. 4C, which shows slices of the 2DIR spectrum taken along the diagonal at which the negative (Δ*A <* 0) signal is maximal. It is difficult to observe this narrowing in the conventional IR spectrum because of the changes in the background absorption upon deformation (whereas in the 2DIR spectrum the background contribution is eliminated because the signal scales nonlinearly with the absorption cross section). After the deformation cycle, the diagonal carbonyl peak has become narrower, mostly due to disappearance of intensity at the high-frequency side. Thus, the narrowing of the hydrogen-bond distribution after a deformation cycle is due to the disappearance of weak hydrogen bonds.

**Figure 4:**
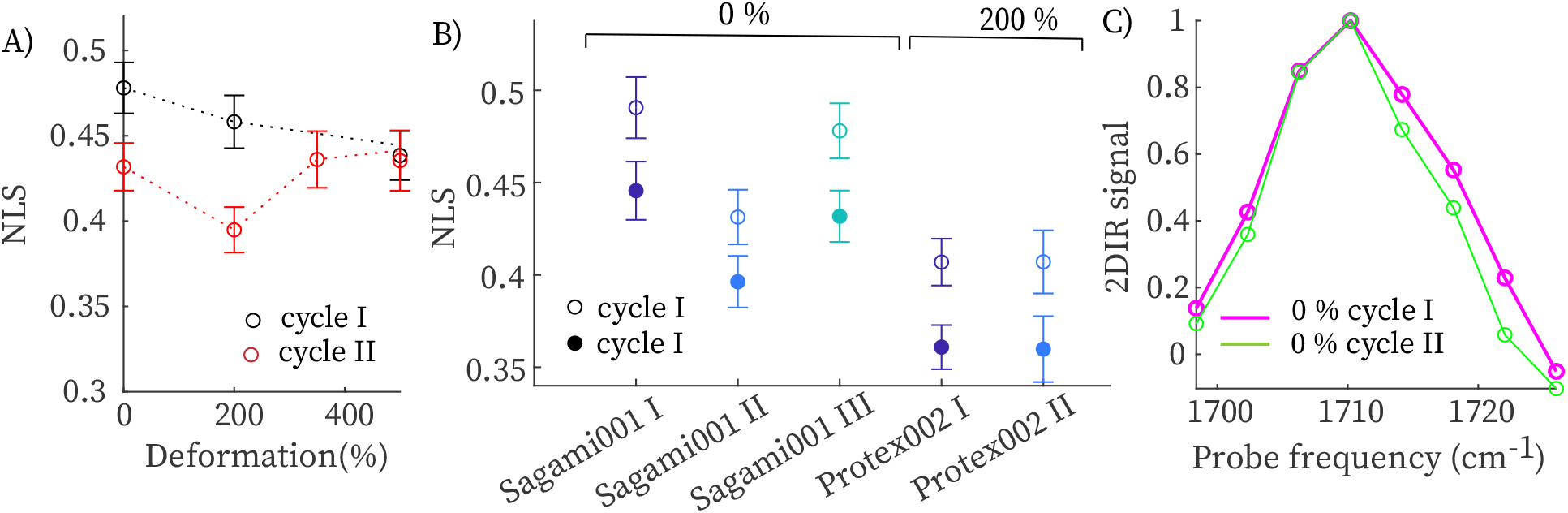
A) Nodal line slope during a deformation cycle. (B) Nodal line slopes before and after a deformation cycle in 5 independent experiments. In the thicker samples (Protex 002) we can only measure 2DIR spectra at 200% strain (for lower deformation the IR absorption is too high). (C) Comparison of bleach diagonal slices extracted by 2DIR spectra in 3B.

The rheo-2DIR results show that a strain cycle causes an irreversible reduction in the number of weak hydrogen bonds in TPU. These weak hydrogen bonds are found mostly in the amorphous regions between the soft and hard domains, where the urethane NH groups form hydrogen bonds mostly to polyether instead of carbonyl groups.^26^ Recent X-ray studies have attributed the strain-softening behavior of TPU to strain-induced softening of these “fuzzy” regions between the hard and soft domains.^1,2^ Our results seem to confirm this idea, and provide a molecular-level explanation of the strain-softening of the fuzzy regions. This picture is confirmed by the lower nodal line slope observed in 2DIR spectra recorded after deforming the sample at high (∼100^*◦*^C) temperature, which probably also destroys the weak hydrogen bonds in the unordered regions (see SI). To conclude, rheo-2DIR spectroscopy sheds new light on the molecular processes that underlie the Mullin effect of polyurethane. Based on these first experiments, we believe that rheo-2DIR can be a valuable addition to the existing physical methods for studying the elastic properties of materials: it can help to improve our understanding of the molecular origin of the elastic response of not only synthetic, but in particular biological polymer-based materials.

## Supporting Information Available

Sample characterization; NLS cycles on Protex 002;Temperature effect on nodal line slope.

