## Supplementary Material for "Rheo-2DIR spectroscopy reveals strain-induced hydrogen-bond redistribution in polyurethane"

### Sample characterization

Samples were obtained by cutting condom in pieces of an area of 1 $cm^2$. To characterize the soft segments, we study the infrared region around 1100 $cm^{-1}$, also called finger-print region. We observe a strong band at 1110 $cm^{-1}$, which is assigned in literature to a stretching mode of $CH_2$-O-$CH_2$ of polyether, suggesting that the soft segments are made by polyether. This is confirmed by the fact we do not observe any strong band at 1180 $cm^{-1}$ that is assigned to C-O-C mode in polyester. For band assignement, see ref.[1]

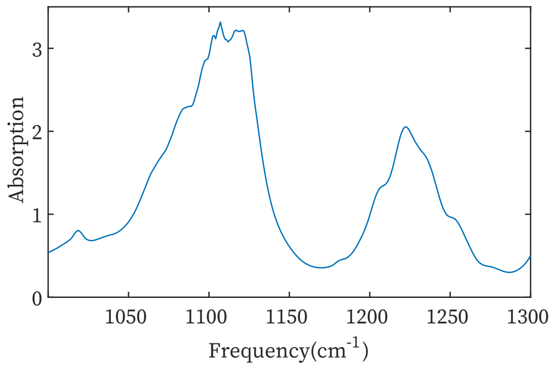


Figure S1: IR spectrum of a sample taken from Sagami 001 at zeros stress condition.

### Nodal line slope Protex 002

We perform rheo-2DIR on sample taken from Protex 002. Because Protex 002 is thicker, the absorption of the carbonyl peaks is higher with respect to the Sagami 001, and thus at zero-stress condition most of the IR light is being absorbed by the sample. Due to this, we can perform rheo-2DIR just when the sample is already deformed, and thus thinner. In Fig.S6 we report the data and analysis of a sample subjected to three consecutive stress cycles up to 600 %. As reported in the main text, we again observe a decrease of the NLS upon recovery of the lowest possible deformation and a shift toward lower frequency of the bleach signal (Fig.S6 C-D).

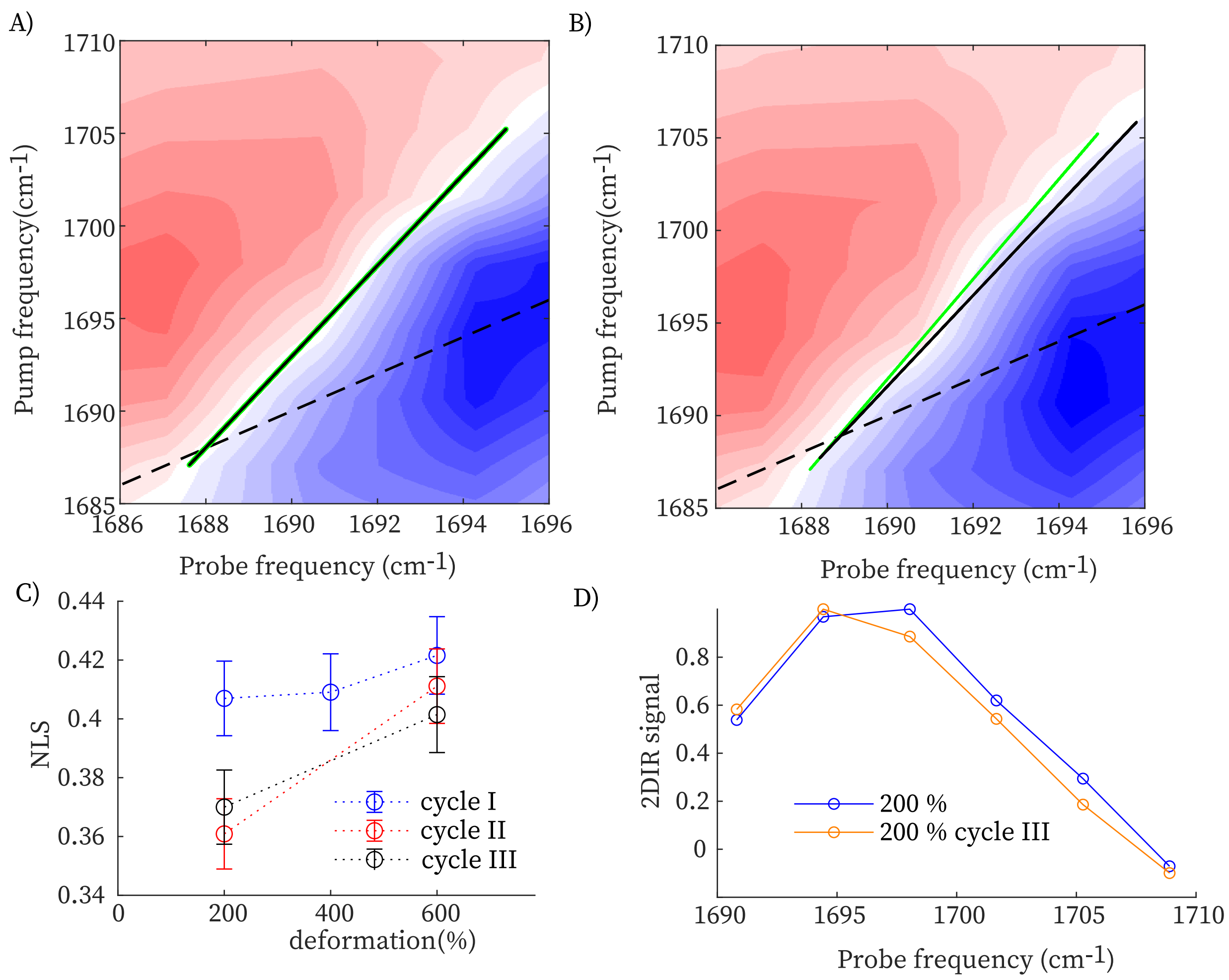


Figure S2: A-B) 2DIR spectra and nodal line slopes of a sample at 200 % of deformation and upon recovery of 200 % stress condition after a deformation up to 600%, respectively. C) NLS slopes for three subsequent cycles. D) Comparison of bleach diagonal slices extracted by 2DIR spectra in A-B.

### Nodal line slope Protex 002 subjected to deformation at high temperature

Samples were deformed at high temperature to reduce the thickness from 30 to 25 $\mu$m. By doing so, we observed that the NLS starts from a lower value with respect to the untreated one, and increases during the cycle.

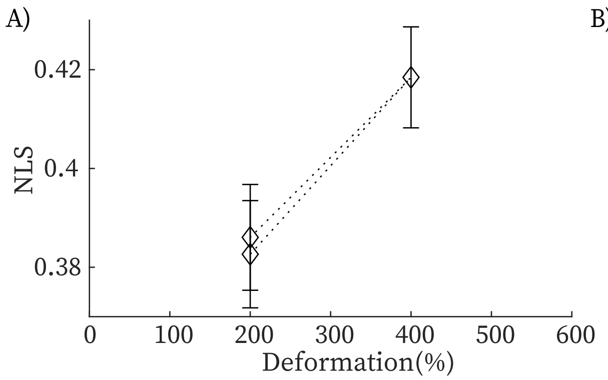


Figure S3: A-B) Nodal line slope of a sample from Protex 002 that was deformed at high temperature (70-100 °C) for two different experiments cycling till 400 % and 600 % respectively.
